# Cognitive functions and jugular venous reflux in severe mitral regurgitation

**DOI:** 10.1101/466037

**Authors:** Shih-Hsien Sung, Ching-Wei Lee, Pei-Ning Wang, Hsiang-Ying Lee, Chen-Huan Chen, Chih-Ping Chung

## Abstract

Cardiac diseases with elevated central venous pressure have higher frequency of jugular venous reflux (JVR), which is associated with decreased cerebral blood flow and white matter hyperintensities. Whether patients with severe mitral-regurgitation (SMR) have poorer cognitive functions and whether JVR is involved were determined. Patients with SMR and age/sex-matched controls were prospectively recruited. Neuropsychological tests such as global cognitive (Mini-Mental State Examination, MMSE), verbal memory, executive, and visuospatial domains were performed. Cardiac parameters by cardiac catheterisation and echocardiography, and the frequency of JVR by colour-coded duplex ultrasonography were obtained. Forty patients with SMR and 40 controls (71.1±12.2, 38–89 years; 75% men) were included. Compared with the controls, patients with SMR had lower scores in all neuropsychological tests but only MMSE and visuospatial test scores were statistically significant after adjusting for age, sex, and educational level. We further adjusted for cardiovascular risk factors; the significance remained in the visuospatial test but diminished in MMSE. Multivariate linear regression analyses adjusted for age, sex, and educational level showed that JVR combined with high right-atrial-pressure (RAP > 50th-percentile, 12 mmHg) was significantly associated with poorer performances in both MMSE [right JVR: B coefficient(95% confidence interval, *p*)=-2.83(−5.46–0.20, 0.036); left JVR: −2.77(−5.52–0.02, 0.048)] and visuospatial test [right JVR: −4.52(−8.89–0.16, 0.043); left JVR: −4.56(−8.81–0.30, 0.037)], with significances that remained after further adjusting for cardiovascular risk factors. Our results suggest that retrogradely-transmitted venous pressure might be involved in the mechanisms mediating the relationship between cardiac diseases and brain functions.

## Introduction

Cerebral venous drainage impairment with elevated venous pressure would decrease cerebral blood flow (CBF), damage the blood–brain barrier (BBB), and lead to brain dysfunctions [1,2]. Internal jugular vein (IJV) is the largest extracranial vein for cerebral venous drainage [1]. Jugular venous reflux (JVR) indicates a retrograde flow in IJV, which usually occurs when the reversed pressure gradient was elevated beyond the capacity of the IJV valves [3]. We previously showed that during Valsalva’s manoeuvre (VM), people with JVR would decrease CBF and dilate retinal venules more than ones without JVR [4,5]. In addition, JVR has been found to be associated with white matter hyperintensities (WMH) in the elderly people [6]. These results indicate that JVR might influence CBF, cerebral microvessels, and brain tissues via elevated venous pressure retrogradely transmitted into the cerebral venous system.

Continuous or repeated elevated venous pressure proximal to IJV might result in wear and tear of the IJV valves and lead to valvular incompetence [3,7,8]. Indeed, certain cardiac diseases with increased central venous pressure such as heart failure or valvular heart disease have a higher frequency of JVR [7,8]. Recently, the number of studies that have reported cognitive impairment in patients with cardiac diseases has been increasing [9]. However, whether cerebral venous return status is a factor involved in the relationship between cardiac diseases and cognitive function remains to be elucidated. The present study compared neuropsychological performances between patients with severe mitral valve regurgitation (SMR) and age- and sex-matched normal controls. Patients with SMR will encounter pulmonary venous hypertension at the beginning, followed by combined pre and post-capillary pulmonary hypertension during disease progression [10,11]. The right ventricular pressure, as well as the right atrial pressure (RAP), increased thereafter, which may lead to IJV valvular incompetence and compromise the cerebral venous return [10,11]. We hypothesised that patients with SMR have poorer cognitive functions and the presence of JVR and/or elevated RAP might be associated with cognitive impairment in these patients.

## Materials and Methods

### Study population

Patients with SMR, referred for surgical intervention in a tertiary medical centre, were eligible for this study. Every patient underwent transthoracic and transoesophageal echocardiography and cardiac catheterisation to confirm the diagnosis and to evaluate the feasibility for surgery. Patients who had disease durations of 1 year or longer from the initial diagnosis to the time of catheterisation and echocardiography were included. Among the eligible patients, those who met the following criteria were excluded from this analysis: (1) had concomitant severe aortic valve disease, mitral stenosis, acute coronary syndrome, or pericardial disease; (2) had unstable haemodynamics, or New York Heart Association functional class IV symptoms; (3) had existing neurological diseases, such as stroke, brain tumour, dementia, or other neurodegenerative diseases; and (4) had significant stenosis (>50%) over the cervical internal carotid and vertebral arteries using neck duplex sonography. Cardiac diseases other than mitral valvular disease were excluded because they might have different mechanisms and effects on the cognitive functions. A total of 40 individuals were included based on these criteria. We also recruited 40 age- and sex-matched normal controls from outpatients who visited our neurological clinics. These normal controls had no cardiac, neurological, or malignant medical histories.

Cardiovascular risk factors were either measured or assessed through self-report. The presence of hypertension was determined by a self-report of current antihypertensive medication prescription or by a measurement of either systolic BP of ≥140 mmHg or diastolic BP ≥90 mmHg [12]. Diabetes mellitus (DM) was defined by either a self-report of current DM medication or a measurement of haemoglobin A1c (HgbA1c) of ≥6.5% [13]. Chronic kidney disease (CKD) was defined according to an estimated glomerular filtration rate (eGFR) of ≤60 mL/min/1.73 m2 [14]. The design of this study was reviewed and approved by the institutional review board of Taipei Veterans General Hospital.

### Cardiac Catheterisation

Cardiac catheterisation was performed in all patients with SMR using a percutaneous approach via the radial artery for coronary angiogram and right IJV for right heart catheterisation. Data of mean pulmonary artery wedge pressure (PAWP), pulmonary artery pressure (PAP), right ventricular pressure (RVP), RAP, mixed venous oxygen saturation (SvO_2_), and cardiac output were obtained. Cardiac output was then divided by body surface area (BSA) to obtain the cardiac index.

### Echocardiography

A comprehensive two-dimensional, M-mode, and Doppler echocardiogram was performed by a skilled echocardiographer using commercially available echocardiographic devices (Philip IE33, Andover, MA, USA) following a standardised protocol. The severity of mitral regurgitation was evaluated according to the AHA/ACC guideline, and an effective regurgitant orifice of ≥0.4 cm^2^ was referred as SMR [15]. Both left and right heart structures and functions were obtained, including left ventricular end-diastolic and end-systolic dimension, left atrial dimension, left atrial volume, and estimated right ventricular systolic pressure. Left ventricular ejection fraction (LVEF) was obtained using biplane Simpson’s method, and left ventricular mass was measured using the area-length method. The peak trans-mitral filling velocity at early diastole (E), septal mitral annulus moving velocity at early diastole (e′), and E/e′ ratio were also obtained. All parameters were measured in triplicate and averaged according to the guideline of the American Society of Echocardiography. Decompensated heart failure was defined as reduced LVEF (<35%) with chronic clinical symptoms (≥6 months) of New York Heart Association functional class III–IV.

### Colour-coded duplex ultrasonography: JVR determination

Neck colour-coded duplex sonography was performed in all patients with SMR using a 7-MHz linear transducer (iU22; Philips, New York, NY, USA) by the same technician who was blinded to subjects’ characteristics. On examination, subjects were in a head-straight, flat supine position after a quiet 10-min rest. The IJV was initially insonated longitudinally and thoroughly from the proximal part of the neck base rostrally to the distal part of the submandibular level to detect any possible spontaneous JVR at baseline. Then, the VM was performed by forcible expiration by the subject via the mouth into a flexible rubber tube connected to a manometer. Subjects were asked to reach the 40 mmHg Valsalva pressure and maintain it for at least 10 s. During the VM, the distal margin window of the colour signal was placed at the tip of the flow divider of the internal carotid artery. The coloured box was adjusted to include the entire lumen of the IJV; if retrograde colour appeared in the centre of the lumen, the retrograde flow would then be confirmed by Doppler spectrum. JVR was determined when the retrograde-flow colour in the centre of the lumen and the Doppler-flow waveform demonstrated reversed flow for >0.5 s spontaneously or/and during VM [3-6].

Routine cervical arterial examination including examination of internal carotid and vertebral arteries was also performed in all patients with SMR.

### Cognitive Function Assessment

All patients with SMR and normal controls underwent a face-to-face neuropsychological examination carried out by trained interviewers. In addition to the global cognitive performance, which was examined using the Mini-Mental State Examination (MMSE), three different cognitive domains (verbal memory, visuospatial function, and executive function) were assessed using extensive neuropsychological tests as follows:

- Verbal memory: delayed (10 min) free recall in the Chinese Version of the Verbal Learning Test (CVVLT) [16].
- Visuospatial function: the copy of the Taylor complex figure test [17].
- Executive function: digit backward test [18].

### Statistical analysis

Analyses were performed using SPSS software (v22.0, IBM, Armonk, NY, USA). All continuous variables are described as mean ± standard deviation (SD) and discrete variables as percentages. Comparisons of case and control were made using non-parametric Mann–Whitney tests. When appropriate, chi-square (χ^2^) or Fisher’s exact tests were performed for categorical variables. Univariate and multivariate linear regression analyses of neuropsychological test scores as the dependent variable were performed. Adjusted confounding factors were age, sex, educational level, and cardiovascular risk factors (hypertension, DM, hyperlipidaemia, cigarette smoking, alcohol consumption, and CKD).

To test our postulation that cerebral venous return status might be involved in the relationship between cognitive impairment and SMR, we analysed the hemodynamic parameters that may affect cerebral venous return, e.g., the RAP and presence of JVR, as independent variables individually. We also used the 50th percentile of the mean RAP, with 12 mmHg as a cut-off point. Three kinds of binary category variables, (1) RAP ≥ and <12 mmHg, (2) the presence or absence of JVR, and (3) the presence or absence of combined JVR and high RAP (≥12 mmHg), were individually analysed as independent variables. Furthermore, since decreased cardiac output is commonly postulated as a contributor to cognitive impairment in cardiac diseases, we also put cardiac index and LVEF into analyses.

## Results

Table 1 shows the demographics and neuropsychological test scores of 40 patients with SMR and 40 age-/sex-matched control. The patient group had higher frequency of cardiovascular risk factors, except cigarette smoking; however, the difference was statistically significant only in the frequency of CKD. Among the patients with SMR, 10 (25%) had decompensated heart failure.

The patient group had lower scores in all neuropsychological tests compared with control group, but only statistically significant in MMSE and Taylor complex figure test and borderline significant in digit backward test after adjusting for age, sex, and educational level. We further adjusted for cardiovascular risk factors, with significance remaining in the Taylor complex figure test (*p* = 0.046) but lower in MMSE (*p* = 0.058).

**Table 1.** Comparisons of demographics and cognitive functions between patients with severe mitral regurgitation and normal controls.

|  | **SMR**<br>**(n = 40)** | **Control**<br>**(n = 40)** | ***P*** |
| --- | --- | --- | --- |
| Age, years, mean (SD, range) | 71.1 (12.2, 38-89) | 71.1 (12.2, 38-89) | - |
| Sex, man, n (%) | 30 (75.0) | 30 (75.0) | - |
| Education, years, mean (SD) | 10.6 (4.8) | 10.3 (4.9) | 0.903 |
| Hypertension, n (%) | 22 (55.0) | 15 (37.5) | 0.178 |
| Diabetes mellitus, n (%) | 8 (20.0) | 5 (12.5) | 0.546 |
| Hyperlipidemia, n (%) | 10 (25.0) | 3 (7.5) | 0.066 |
| Cigarette smoking, n (%) | 12 (30.0) | 13 (32.5) | 1.000 |
| Chronic kidney disease, n (%) | 20 (50.0) | 5 (5.0) | <0.001 |
| **Age, sex, education adjusted** |  |  |  |
| MMSE, mean (SD) | 26.1 (5.1) | 27.8 (2.4) | 0.020 |
| Verbal memory: CVVLT 10 min, mean (SD) | 6.4 (3.0) | 6.9 (1.8) | 0.387 |
| Executive function: digit backward test, mean (SD) | 5.4 (2.7) | 6.4 (3.0) | 0.081 |
| Visuospatial function: the Taylor complex figure test, mean (SD) | 29.8 (6.9) | 31.9 (4.0) | 0.040 |

SMR = severe mitral regurgitation; CVVLT = Chinese Version of the Verbal Learning Test.

Table 2 shows the hemodynamic parameters measured by cardiac catheterization and the frequency of JVR detected by color-coded duplex ultrasonography in SMR patients. An elevated mean RAP and high frequency of JVR were observed in patients with SMR.

**Table 2.** Hemodynamic parameters in patients with severe mitral regurgitation.

|  |  |
| --- | --- |
| LV ejection fraction, %, (SD, range) | 47.3 (12.6, 25-70) |
| Cardiac index, l/min/m<sup>2</sup>, (SD, range) | 3.5 (1.0, 2.1-6.1) |
| PAWP, mmHg, (SD, range) | 25.2 (10.4, 9-47) |
| PAP, mmHg, (SD, range) | 34.7 (10.7, 17-59) |
| RVP, mmHg, (SD, range) | 12.3 (7.3, 1-34) |
| RAP, mmHg, (SD, range) | 11.9 (5.9, 4-24) |
| Right JVR, n (%) | 20 (50.0) |
| Left JVR, n (%) | 22 (55.0) |

LV = left ventricle; PAWP = pulmonary artery wedge pressure; PAP = pulmonary artery pressure;
RVP = right ventricular pressure; RAP = right atrial pressure; JVR = jugular venous reflux.

We then performed multivariate analyses to test which haemodynamic parameter was associated with poorer cognitive domains, e.g., MMSE and the Taylor complex figure test, in patients with SMR (Table 3). Multivariate analyses adjusted for age, sex, and educational level showed that cardiac index, LVEF, mean RAP, high mean RAP (≥12 mmHg), or presence of right or left JVR were not associated with the MMSE and Taylor complex figure test scores. However, JVR combined with high mean RAP was significantly associated with poorer performances both in MMSE and Taylor complex figure test. The significances remained after further adjusting for cardiovascular risk factors. We also divided patients into four groups according to the presence of absence of JVR and high mean RAP. Fig 1 shows the mean scores of MMSE and Taylor complex figure test of the four groups. Cognitive functions in patients with isolated JVR or high mean RAP were not poorer than those with the absence of JVR and high mean RAP; however, JVR combined with high mean RAP had the lowest scores in both MMSE and Taylor complex figure test among the four groups. Multivariate analyses showed that patients in the group of JVR combined with high mean RAP had significantly poorer performances in both MMSE and Taylor complex figure test compared with those in the other three groups.

**Figure 1.**
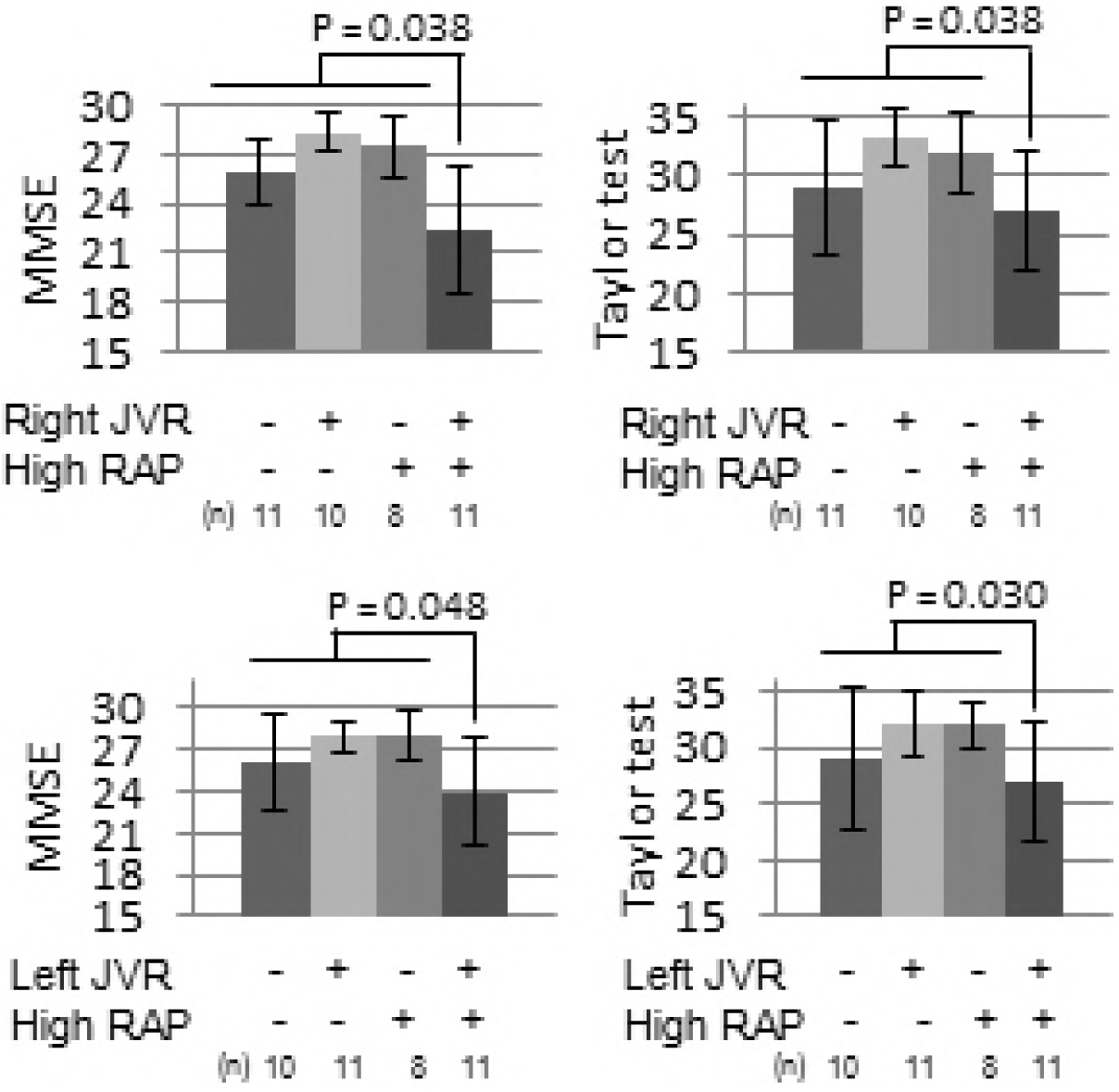
Cognitive functions in four groups of patients with severe mitral valve regurgitation classified according to the presence or absence of jugular venous reflux and high right atrial pressure.

**Table 3.** Associations of cardiac parameters with cognitive functions in patients with severe mitral regurgitation.

|  | **Mini-Mental Status Examination** |  |  |  |
| --- | --- | --- | --- | --- |
|  | B (95% CI) | *P*<sup>a</sup> | B (95% CI) | *P*<sup>b</sup> |
| LVEF | 0.04 (-0.08-0.16) | 0.532 |  |  |
| Cardiac index | -1.03 (-2.77-0.72) | 0.232 |  |  |
| RAP | -0.05 (-0.27-0.17) | 0.640 |  |  |
| RAP > 12 mmHg | -1.33 (-3.79-1.13) | 0.278 |  |  |
| Right JVR | -1.14 (-4.49-2.21) | 0.494 |  |  |
| Left JVR | -0.72 (-4.04-2.61) | 0.664 |  |  |
| Right JVR & RAP > 12<br>mmHg | -2.83 (-5.46-0.20) | 0.036 | -3.05 (-5.92-0.19) | 0.038 |
| Left JVR & RAP > 12 mmHg | -2.77 (-5.52-0.02) | 0.048 | -2.96 (-5.89-0.02) | 0.048 |
|  | **Visuospatial function: the Taylor complex figure test** |  |  |  |
|  | B (95% CI) | *P*<sup>a</sup> | B (95% CI) | *P*<sup>b</sup> |
| LVEF | 0.10 (-0.05-0.28) | 0.237 |  |  |
| Cardiac index | 0.10 (-2.58-2.79) | 0.936 |  |  |
| RAP | -0.08 (-0.44-0.28) | 0.648 |  |  |
| RAP > 12 mmHg | -0.92 (-4.93-3.10) | 0.642 |  |  |
| Right JVR | -1.51 (-6.11-3.09) | 0.507 |  |  |
| Left JVR | -3.05 (-7.47-1.37) | 0.169 |  |  |
| Right JVR & RAP > 12<br>mmHg | -4.52 (-8.89-0.16) | 0.043 | -4.93 (-9.56-0.30) | 0.038 |
| Left JVR & RAP > 12 mmHg | -4.56 (-8.81-0.30) | 0.037 | -4.96 (-9.40-0.52) | 0.030 |

LVEF = left ventricle ejection fraction; RAP = right atrial pressure; JVR = jugular venous reflux; B = B coefficient; CI = confidence interval.
<sup>a</sup>adjusted for age, sex and education years. <sup>b</sup>adjusted for age, sex, education years and cardiovascular risk factors (hypertension, diabetes mellitus, hyperlipidemia, cigarette smoking, alcohol consumption, and chronic kidney disease).

## Discussion

The main findings were that patients with SMR had (1) poorer global cognitive (MMSE) and visuospatial (the Taylor figure test) functions compared with those in normal controls and (2) JVR combined with high RAP was associated with these cognitive impairments.

We previously reported that the prevalence of JVR in the general population (16–89 years old) is approximately 18–36% on the right side and 6–29% on the left side [19]. The present study showed a high frequency of JVR (50–55%) in patients with SMR. Chronic SMR with a continuous or repeated elevated central venous pressure might wear and tear the IJV valves and lead to valvular incompetence. This postulation is supported by a high RAP found in our SMR patients and the other studies showing a higher frequency of JVR in heart failure or tricuspid valve disease which have elevated central venous via elevated RAP.

Although retrogradely transmitted venous pressure by JVR has been shown to reach the cerebral venous system and influence CBF [3-6], the extent of induced cerebral venous hypertension is milder than that of the other conditions, such as dural arteriovenous fistula (DAVF) [20-22]. Therefore, compared with diffuse cerebral white matter hyperintensities (WMH) caused by DAVF [20-22], JVR is only associated with WMH over caudal brain (occipital, thalamus, and infratentorial brain regions) in which venous drainage pathway is closer to IJV [6]. In addition, age is needed to enhance JVR-related brain insults; JVR is associated with the severity of WMH only in people aged ≥75 years [6]. The present study had similar observations. Merely the presence of JVR was not associated with SMR-related cognitive impairment; nevertheless, with the additional high RAP, JVR was associated with poorer cognitive performances, global cognitive (MMSE), and visuospatial (the Taylor figure test) functions in patients with SMR (Fig 1). Our results lead to the postulation that high RAP related to heart failure has limited influence on the brain if IJV valves are competent; JVR with high RAP can cause brain dysfunction via retrogradely transmitted venous pressure only when IJV valves are incompetent (Fig 2).

**Figure 2.**
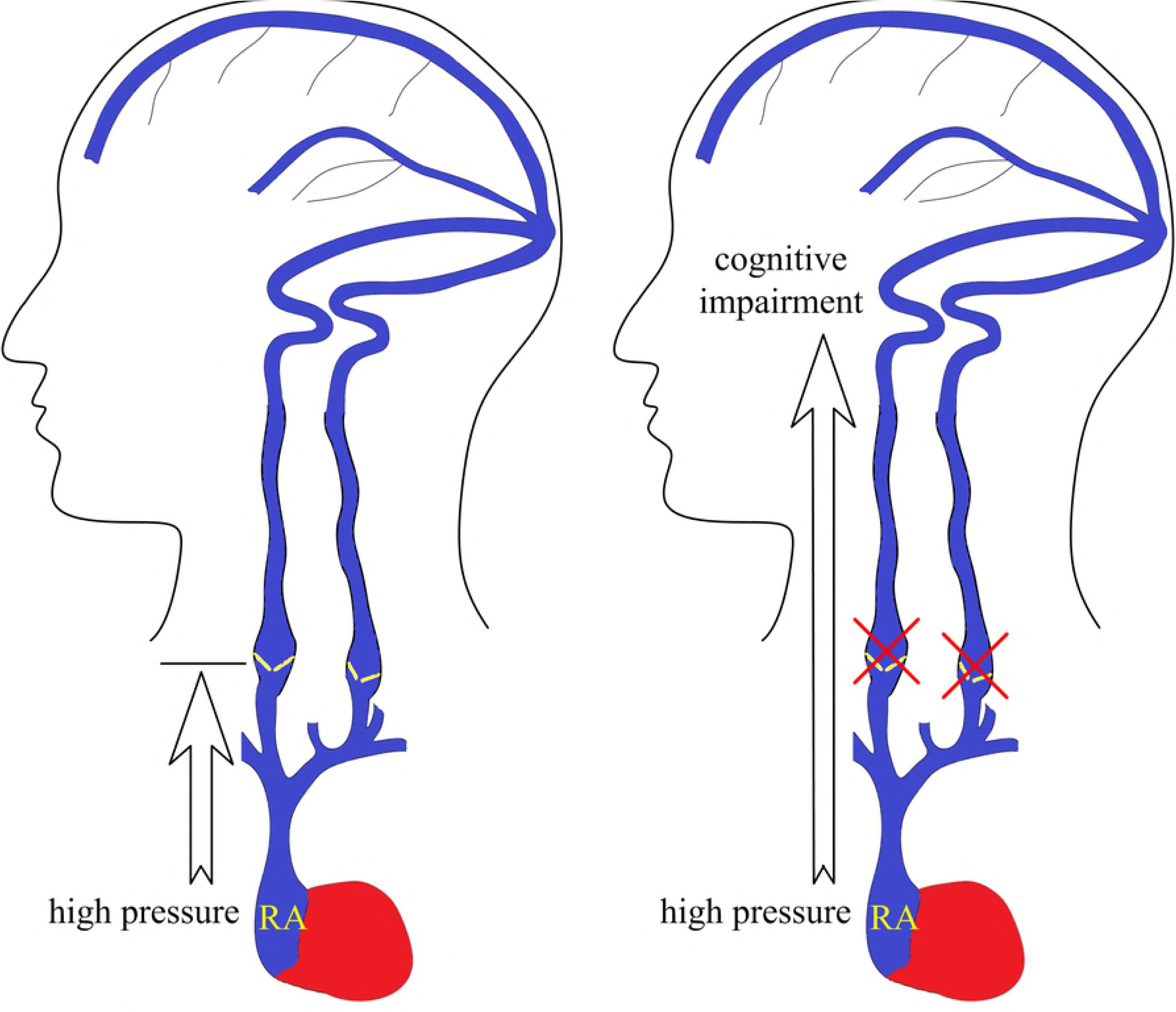
Postulated role of jugular venous valve incompetence (jugular venous reflux) in the mechanisms mediating the relationship between cardiac diseases with elevated right atrial pressure and cognitive impairment.

Several studies on brain–heart axis have emerged, and they have shown a relationship between cognitive impairment and cardiac diseases [9]. Most studies were focusing on heart failure and little on the effect of mitral valve disease on cognitive functions [9,23]. Our results showed that compared with age- and sex-matched normal controls, patients with SMR had poorer global cognitive performance (MMSE) and visuospatial function (Taylor figure test) after adjusting with educational level. The diminished significance of association in MMSE after adjusting for cardiovascular risk factors suggests that more prevalent cardiovascular risk factors such as hypertension, DM, and CKD might be contributors to poorer global cognitive function in SMR. Notably, the anatomic correlations of visuospatial function impairment, significantly and independently associated with SMR, include the occipital lobe, which is one of the JVR-susceptible regions [6]. This result also supports our postulated mechanism mediating the cognitive impairment in SMR (Fig 2).

Cerebral circulation includes artery supply and venous drainage. Both of them are responsible for adequate CBF and brain metabolic homeostasis [2,24]. Recently, several studies have indicated that, in addition to maintaining adequate CBF and BBB function, waste and lymphatic clearance are dependent on cerebral venous drainage [25-27]. However, a greater proportion of studies are focusing on the arterial side, e.g., cardiac output, when evaluating the relationship between the circulation (heart) and the brain [9]. Results of the present study indicate a role of the venous side in the impact of cardiac disease on brain dysfunction. We did not find associations between parameters reflecting the arterial side, such as cardiac index and LVEF and cognitive functions in patients with SMR. Our results are consistent with those of a recent study [28]. They investigated the association between various cardiac haemodynamic parameters and the volume of WMH in chronic valvular heart disease such as mitral valve regurgitation (43.1% of the study population) and found that RAP is associated with WMH. In their results, instead of cardiac index, LVEF, and other cardiac hemodynamic parameters, only the mean RAP is significantly, independently, and linearly associated with the WMH volume. However, they did not investigate the neurological functions and competence of IJV valves in those patients. The role of JVR on these valvular heart disease-related WMHs and whether WMH is associated with cognitive impairment as shown in our study were unclear.

The present study has limitations. The study sample size was relatively small. In addition, the cross-sectional study setting could not establish a causal relationship. Therefore, a larger and longitudinal study is necessary to validate our postulation. In addition, more investigated tools such as brain imaging are needed to further evaluate the underlying mechanisms between the cerebral venous drainage impairment and cognitive abnormalities in SMR.

## Conclusions

Patients with SMR had poorer cognitive function, particularly in the visuospatial domain, and JVR combined with high RAP was associated with poorer visuospatial function in these patients. The results suggest that retrogradely transmitted venous pressure but not low cardiac output might be involved in the mechanisms mediating the relationship between valvular heart disease and brain functions. In addition to management for decreasing RAP, IJV valve repair might be a potential treatment option for cardiac disease-related brain dysfunctions.

## Acknowledgements

The authors received grants from Ministry of Science and Technology and Taipei Veterans General Hospital, Taiwan (Chung: VGH V105C-055; MOST 104-2314-B-075-MY3; Wang: NSC 101-2314-B-010; NSC 102-2314-B-010-051-MY2; Taipei VGH V104C-059).

## References

1. Schaller B. Physiology of cerebral venous blood flow: from experimental data in animals to normal function in humans. Brain Research Reviews 2004;46:243–260.

2. Schaller B, Graf R. Cerebral venous infarction: the pathophysiological concept. Cerebrovasc Dis 2004;18:179–188.

3. Chung CP, Hu HH. Jugular venous reflux. Journal of Medical Ultrasound 2008;16:210–222.

4. Wu IH, Sheng WY, Hu HH, Chung CP. Jugular venous reflux could influence cerebral blood flow: a transcranial Doppler study. Acta Neurol Taiwan 2011;20:15–21.

5. Chung CP, Hsu HY, Chao AC, Cheng CY, Lin SJ, Hu HH. Jugular venous reflux affects ocular venous system in transient monocular blindness. Cerebrovasc Dis 2010;29:122–129.

6. Chung CP, Wang PN, Wu YH, Tsao YC, Sheng WY, Lin KN, et al. More severe white matter changes in the elderly with jugular venous reflux. Ann Neurol 2011;69:553–559.

7. Fisher J, Vaghaiwalla F, Tsitlik J, Levin H, Brinker J, Weisfeldt M, et al. Determinants and clinical significance of jugular venous valve competence. Circulation 1982;65:188–196.

8. Dresser LP, McKinney WM. Anatomic and pathophysiologic studies of the human internal jugular valve. Am J Surg 1987;154:220–224.

9. Cannon JA, Moffitt P, Perez-Moreno AC, Walters MR, Broomfield NM, McMurray JJV, et al. Cognitive Impairment and Heart Failure: Systematic Review and Meta-Analysis. J Card Fail 2017;23:464–475.

10. Asgar AW, Mack MJ, Stone GW. Secondary mitral regurgitation in heart failure: pathophysiology, prognosis, and therapeutic considerations. J Am Coll Cardiol 2015;65:1231–1248.

11. Harb SC, Griffin BP. Mitral Valve Disease: a Comprehensive Review. Curr Cardiol Rep 2017;19:73.

12. Jones DW, Hall JE. Seventh report of the Joint National Committee on Prevention, Detection, Evaluation, and Treatment of High Blood Pressure and evidence from new hypertension trials. Hypertension 2004;43:1–3.

13. American Diabetes Association. Diagnosis and classification of diabetes mellitus. Diabetes Care 2010;33 Suppl 1:S62–69.

14. Levey AS, Eckardt KU, Tsukamoto Y, Levin A, Coresh J, Rossert J. Definition and classification of chronic kidney disease: a position statement from Kidney Disease: Improving Global Outcomes (KDIGO). Kidney Int 2005;67:2089–2100.

15. Nishimura RA, Otto CM, Bonow RO, Carabello BA, Erwin JP 3rd, Fleisher LA, et al. 2017 AHA/ACC Focused Update of the 2014 AHA/ACC Guideline for the Management of Patients With Valvular Heart Disease: A Report of the American College of Cardiology/American Heart Association Task Force on Clinical Practice Guidelines. J Am Coll Cardiol 2017;70:252–289.

16. Chang CC, Kramer JH, Lin KN, Chang WN, Wang YL, Huang CW, et al. Validating the Chinese version of the Verbal Learning Test for screening Alzheimer’s disease. Journal of the International Neuropsychological Society: JINS 2010;16:244–251.

17. Liu HC, Lin KN, Teng EL, Wang SJ, Fuh JL, Guo NW, et al. Prevalence and subtypes of dementia in Taiwan: a community survey of 5297 individuals. Journal of the American Geriatrics Society 1995;43:144–149.

18. Hester RL, Kinsella GJ, Ong B. Effect of age on forward and backward span tasks. Journal of the International Neuropsychological Society: JINS 2004;10:475–481.

19. Chung CP, Lin YJ, Chao AC, Lin SJ, Chen YY, Wang YJ, et al. Jugular venous hemodynamic changes with aging. Ultrasound Med Biol 2010;36:1776–1782.

20. Waragai M, Takeuchi H, Fukushima T, Haisa T, Yonemitsu T. MRI and SPECT studies of dural arteriovenous fistulas presenting as pure progressive dementia with leukoencephalopathy: a cause of treatable dementia. Eur J Neurol 2006;13:754–759.

21. Yamakami I, Kobayashi E, Yamaura A. Diffuse white matter changes caused by dural arteriovenous fistula. J Clin Neurosci 2001;8:471–475.

22. Zeidman SM, Monsein LH, Arosarena O, Aletich V, Biafore JA, Dawson RC, et al. Reversibility of white matter changes and dementia after treatment of dural fistulas. AJNR Am J Neuroradiol 1995;16:1080–1083.

23. Nikendei C, Schäfer H, Weisbrod M, Huber J, Geis N, Katus HA, et al. The Effects of Mitral Valve Repair on Memory Performance, Executive Function, and Psychological Measures in Patients With Heart Failure. Psychosom Med 2016;78:432–442.

24. Pantoni L. Cerebral small vessel disease: from pathogenesis and clinical characteristics to therapeutic challenges. Lancet Neurol 2010;9:689–701.

25. Kress BT, Iliff JJ, Xia M, Wang M, Wei HS, Zeppenfeld D, et al. Impairment of paravascular clearance pathways in the aging brain. Ann Neurol 2014;76:845–861.

26. Iliff JJ, Wang M, Liao Y, Plogg BA, Peng W, Gundersen GA, et al. A paravascular pathway facilitates CSF flow through the brain parenchyma and the clearance of interstitial solutes, including amyloid β. Sci Transl Med 2012;4:147ra111.

27. Louveau A, Smirnov I, Keyes TJ, Eccles JD, Rouhani SJ, Peske JD, et al. Structural and functional features of central nervous system lymphatic vessels. Nature 2015;523:337–341.

28. Lee WJ, Jung KH, Ryu YJ, Kim JM, Lee ST, Chu K, et al. Association of Cardiac Hemodynamic Factors With Severity of White Matter Hyperintensities in Chronic Valvular Heart Disease. JAMA Neurol 2018;75:80–87.

